# Outcrossing increases resistance against coevolving parasites

**DOI:** 10.1101/2024.01.30.578011

**Authors:** Samuel P. Slowinski, Jennifer D. Gresham, McKenna J. Penley, Curtis M. Lively, Levi T. Morran

## Abstract

Despite substantial costs, biparental sex is the dominant mode of reproduction across plant and animal taxa. The Red Queen hypothesis (RQH) posits that coevolutionary interactions with parasites can favor biparental sex in hosts, despite the costs. In support of the RQH, previous studies found that coevolutionary interactions with virulent bacterial parasites maintained high outcrossing rates in populations of the androdioecious nematode host *Caenorhabditis elegans*. Here we test three non-mutually exclusive mechanisms that could explain how coevolving parasites maintain outcrossing rates in *C. elegans* hosts: 1) short-term parasite exposure induces plastic increases in the hosts’ propensity to outcross, 2) hosts evolve increased outcrossing propensity in response to selection imposed by coevolving parasites, and 3) outcrossed offspring incur less parasite-mediated fitness loss than selfed offspring, increasing host male frequencies and opportunities for outcrossing. We find no evidence that parasites cause plastic or evolved changes in host outcrossing propensity. However, parental outcrossing significantly increases survival of host offspring in the F2 generation when exposed to a coevolving parasite. Hence, coevolving parasites maintain outcrossing in host populations by selecting against selfed offspring, rather than by inducing changes in the propensity to outcross.

## Introduction

Despite substantial costs (Maynard Smith 1978, Gibson et al. 2017), biparental sex is the dominant mode of reproduction across most plant and animal taxa (reviewed in Bell 1982, Vrijenhoek 1998). The Red Queen hypothesis (RQH) posits that coevolutionary interactions with parasites can produce negative frequency-dependent selection on hosts, perhaps favoring biparental sexual reproduction in host populations despite the costs (Jaenike 1978, Hamilton 1980). The RQH has received substantial empirical support in studies of natural populations (Lively 1987, 1992, Jokela and Lively 1995, Lively and Jokela 1996, Dybdahl and Lively 1998, Krist et al. 2000, Lively and Dybdahl 2000, Lively and Jokela 2002, Duncan and Little 2007, Jokela et al. 2009, King and Lively 2009, Wolinska and Spaak 2009, Ellison et al. 2011, King et al. 2011, Verhoeven and Biere 2013). Additionally, several recent experimental evolution studies also supported the RQH by demonstrating that coevolutionary interactions with the virulent bacterial parasite *Serratia marcescens* can maintain high outcrossing rates in the androdioecious nematode host *Caenorhabditis elegans* (Morran et al. 2011, Slowinski et al. 2016, Slowinski et al. 2023). *C. elegans* populations are composed of males and hermaphrodites. Hermaphrodites are capable of reproducing by self-fertilization (uniparental sex) or by mating with males and outcrossing their eggs with male sperm (biparental sex). The mechanisms by which coevolving parasites select for and maintain high outcrossing rates in *C. elegans* host populations remain unknown.

In the present study we test three non-mutually exclusive mechanisms by which coevolutionary interactions with parasites could maintain high outcrossing rates in *C. elegans* host populations. First, we propose that *C. elegans* could plastically increase their outcrossing propensity in response to exposure to parasites (plasticity-of-outcrossing-propensity hypothesis). We measure outcrossing propensity as the frequency with which *C. elegans* hermaphrodites will outcross, given a standardized opportunity for outcrossing with a male. Second, we propose that host populations could evolve an increased propensity to outcross in response to prolonged selection (across many host generations) from coevolving parasites (evolution-of-outcrossing-propensity hypothesis). Third, we propose that selfed offspring experience higher mortality relative to outcrossed offspring in an environment with coevolving parasites (selection-against-selfed-offspring hypothesis). Because almost all selfed offspring are hermaphrodites, whereas outcrossing produces a 1/1 ratio of males/hermaphrodites (Brenner 1974), selective culling of selfed offspring by the parasite would result in the surviving host population being enriched for males. The resulting increase in male frequencies would increase the opportunity for outcrossing in host populations coevolving with the parasite and could maintain high outcrossing rates in host populations, even in the absence of parasite-induced changes in outcrossing propensity.

### Plasticity-of-outcrossing-propensity hypothesis

The RQH predicts that hosts should be selected to diversify their offspring when exposed to parasites, because coevolving parasites are selected to infect common host genotypes (Jaenike 1978, Hamilton 1980). In support of this prediction, previous studies have demonstrated that some hosts can plastically (within one generation) diversify their offspring in response to parasite exposure. For example, Singh et al. (2015) demonstrated that *Drosophila melanogaster* increases the production of recombinant offspring following infection with parasites. Soper et al. (2014) showed that both males and females in the freshwater snail host *Potamopyrgus antipodarum* increased their number of different mating partners when exposed to a sterilizing trematode parasite. Furthermore, Hite et al. (2017) demonstrated that female *Daphnia dentifera* hosts experimentally infected with the virulent fungal parasite *Metschnikowia bicuspidata* allocated more resources than uninfected control hosts to male offspring production (at the expense of allocating resources into the production of asexual females). These responses to parasite exposure could benefit the host by increasing the production of rare (and presumably resistant) offspring genotypes.

We propose that *C. elegans* could also plastically diversify their offspring in response to parasite exposure, by increasing their propensity to reproduce by outcrossing as opposed to self-fertilization whenever a parasite is detected in the environment. This “plasticity-of-outcrossing-propensity hypothesis” predicts that *C. elegans* that experience a short-term (within one generation) parasite exposure should exhibit a higher propensity to outcross relative to unexposed *C. elegans*. This hypothesis already has some support in *C. elegans*; Wu et al. (2023) reported that wildtype *C. elegans* hermaphrodites exposed briefly to the pathogenic bacteria *Pseudomonas aeruginosa* outcrossed more relative to hermaphrodites exposed only to an avirulent bacterial food source (*Escherichia coli*). However, the ability of our model parasite, *S. marcescens*, to induce plastic increases in *C. elegans* outcrossing propensity has not been previously tested. Hence it is unknown whether parasite-induced plastic increases in host outcrossing propensity could help explain the observation that continual exposure to coevolving *S. marcescens* parasites maintains high outcrossing rates in *C. elegans* populations (Morran et al. 2011, Slowinski et al. 2016, Slowinski et al. 2023). To test the plasticity-of-outcrossing-propensity hypothesis, we compared the outcrossing propensity (in a parasite-free environment) of *C. elegans* that had recently experienced a brief exposure to the virulent bacterial parasite *S. marcescens* (parasite-exposure treatment) versus the outcrossing propensity of *C. elegans* that had been maintained continuously on their avirulent *E. coli* food source (no-parasite-control treatment).

### Evolution-of-outcrossing-propensity hypothesis

Our second proposed mechanism to explain how coevolving parasites maintain high outcrossing rates in *C. elegans* host populations is that host populations that coevolve with a virulent parasite evolve a higher propensity to reproduce by outcrossing. We test this prediction by assaying the outcrossing propensity (in a parasite-free environment) of *C. elegans* populations that were passaged with a virulent *S. marcescens* parasite strain (SM2170) in a previous study (Slowinski et al. 2023) under three different parasite treatments for 24 host generations prior to our assays. In the heat-killed parasite treatment (1), *C. elegans* hosts were passaged in an environment with heat-killed (avirulent) *S. marcescens* parasites. In the fixed-genotype parasite treatment (2), *C. elegans* hosts were passaged with a fixed-genotype (non-evolving) virulent strain of the *S. marcescens* parasite. In the copassaged parasite treatment (3), *C. elegans* hosts were copassaged with a virulent strain of *S. marcescens*. In the copassaged treatment, both host and parasite were permitted to evolve and potentially to coevolve. In the previous study, host populations in the copassaged treatment evolved and maintained higher outcrossing rates than host populations in the heat-killed parasite or fixed parasite treatments (Slowinski et al. 2023). In the present study, following 24 generations of experimental evolution, we removed host populations from their parasite-treatment environments and assayed their outcrossing propensity in a parasite-free environment to isolate the effects of each host population’s history of (co)evolutionary interactions with parasites on outcrossing propensity.

### Selection-against-selfed-offspring hypothesis

We hypothesize that outcrossing will increase offspring resistance because coevolving parasites are selected to target common host genotypes, and outcrossing can break up common host genotypes and produce novel or rare genotypic variants in the offspring generation. Furthermore, outcrossing reduces similarity between parents and their progeny, which could enhance offspring resistance if coevolving parasites are adapted to infect parental genotypes (Clay and Kover 1996, Keller and Waller 2002). Therefore, we predict that selfed host offspring should exhibit higher mortality than outcrossed host offspring when exposed to coevolving parasites. Consistent with this expectation, several studies have found that susceptibility to infectious parasites is higher in hosts that are inbred (Acevedo-Whitehouse et al. 2003, Spielman et al. 2004) or selfed (Ellison et al. 2011, Masri et al. 2013) (reviewed in King and Lively 2012), and that host genetic variation and outcrossing can facilitate adaptation to a novel parasite (Parrish et al. 2016).

Because half of outcrossed offspring are males while almost all selfed offspring are hermaphrodites (Brenner 1974), if selfed offspring experience higher mortality on the parasite then parasite exposure should result in a higher frequency of males in the surviving host population. Higher male frequencies provide more opportunity for outcrossing, which in turn can lead to higher outcrossing rates (Morran et al. 2009). Therefore, selective culling by parasites of selfed offspring (primarily hermaphrodites) could result in the maintenance of higher host population male frequencies, and consequently higher outcrossing rates, even in the absence of any plastic or evolved changes in the propensity of hosts to reproduce by outcrossing. We test this proposed mechanism by comparing the survival of selfed versus outcrossed *C. elegans* offspring exposed to the virulent coevolved bacterial parasite *S. marcescens*, using host and parasite populations that had been copassaged for 24 host generations (Slowinski et al. 2023). We predict that selfed offspring should exhibit higher mortality than outcrossed offspring when experimentally exposed to the parasite.

## Methods

### Bacterial and nematode strains

For all experiments involving the *Serratia marcescens* parasite, the virulent strain Sm2170 was used. The strain Sm2170 was obtained from S. Katz at Rogers State University (Claremore, OK). All experimentally evolved parasite lines used in assays were established by copassaging Sm2170 with *C. elegans* in a previous study (Slowinski et al. 2023). The avirulent *Escherichia coli* strain OP50 was used as a food source for the nematode hosts in all experiments.

The nematode host strains N2 and CB4856 were obtained from the Caenorhabditis Genetics Center. The genetically diverse host strains CW1-30 and CF3 were established by inbreeding and mutagenizing CB4856 and then passaging it in a neutral (parasite-free) environment. All host lines with a history of (co)evolution with the *S. marcescens* parasite were established by experimentally passaging CF3 (the experimental evolution ancestor) in a previous study (Slowinski et al. 2023).

More detailed description of the source of the host and parasite lines used is available in the supplementary material, in Supplemental Figure 1, and in (Slowinski et al. 2023).

### Plasticity-of-outcrossing-propensity assay 1

The plasticity of outcrossing propensity was measured in two different assays. In both assays, *C. elegans* populations were synchronized (see supplementary material for detailed protocol) and grown at 20 °C on nematode growth media (NGM) petri plates seeded with *E. coli*. In assay 1, when worms reached the fourth stage of larval development (L4), hermaphrodites were exposed to either the *S. marcescens* parasite strain SM2170 (parasite-exposure treatment) or to the *E. coli* food source (control-exposure treatment) for two hours. Following the exposure treatment, one randomly selected hermaphrodite and one randomly selected male (males were unexposed in both treatments) were picked onto each mating plate, based on the protocol described in (Bahrami and Zhang 2013), which we will refer to as the two-worm protocol. Mating plates are small NGM plates that serve as arenas in which replicate hermaphrodites can experience a standardized opportunity to reproduce by outcrossing with a male or by self-fertilization.

Two-worm protocol mating plates were constructed by pouring 4 mL of autoclaved NGM Lite (US Biological, Swampscott, MA) into a 4 cm Petri dish. The mating plates were seeded with 50 μl of *E. coli*, which was dropped on the surface of the agar in the middle of the Petri dish and not spread. All mating plates were incubated at 28 °C for 24 hours to allow the *E. coli* to grow prior to transferring nematodes onto the mating plates. This process was repeated to create replicate mating plates for each experimental nematode line and treatment that we assayed.

We will refer to the individuals that we picked onto the mating plates as the “parents,” because they are the reproductive individuals whose outcrossing propensity we assayed. The parents were left together on mating plates at 20 °C for 48 hours during which time the hermaphrodite on each plate had the opportunity to mate with the male and reproduce by outcrossing, or, alternatively, to reproduce by self-fertilization. After 48 hours, the parents, which had matured into adults, were removed from the mating plates, and their offspring, which were mostly eggs or early larval stages, and which were easily distinguishable from the parents, were left behind to mature on the mating plates. 48 hours after the parents were removed, we scored all of the offspring (which had matured into adults) on each mating plate as either a hermaphrodite/female or a male.

In assay 1, the outcrossing propensity of the inbred lab strain N2 and of the genetically diverse strain CW1-30 (derived from lab strain CB4856) was measured. Unexposed males from the obligately outcrossing line F5 were the source of the males for both treatments.

### Plasticity-of-outcrossing-propensity assay 2

In the plasticity-of-outcrossing-propensity assay 2, the outcrossing propensity of 10 hermaphrodites was measured simultaneously on each mating plate, based on the protocol described in (Wu et al. 2023), which we will refer to as the 20-worm protocol. Assay 2 mating plates were constructed by pouring 10 mL of NGM lite into a 6 cm Petri dish and were seeded by dropping 100 μl of *E. coli* into the center of the plate and then spreading it. Worms were synchronized and grown until the adult stage. When hermaphrodites were first starting to lay eggs, they were exposed to the parasite or *E. coli* control for four hours. Following exposure, 10 randomly selected hermaphrodites were picked onto each mating plate with 10 randomly selected (unexposed) males, and left together for four hours, during which time they had the opportunity to mate. Following this mating opportunity, each hermaphrodite was picked off of the mating plate onto her own plate where she could lay eggs. The number of male and hermaphrodite offspring on each plate was counted three days later.

The following *C. elegans* lines were used in assay 2: the inbred lab strains N2 and CB4856, the genetically diverse strain CF3 (which served as the ancestor for experimental evolution in (Slowinski et al. 2023)), and two hermaphrodite lineages that were isolated from populations that had been experimentally copassaged with *S. marcescens* for 24 host generations (Slowinski et al. 2023). Unexposed males from a genetically matched obligately outcrossing (fixed for *fog-2(q71)* population were used in each assay.

### Evolution-of-outcrossing-propensity assay

In the evolution-of-outcrossing-propensity assay, the outcrossing propensity of *C. elegans* populations that had previously evolved (in (Slowinski et al. 2023)) for 24 generations on either a heat-killed parasite, a fixed-genotype parasite, or a copassaged parasite treatment was measured using the two-worm protocol. Outcrossing propensity was measured as described for the plasticity-of-outcrossing-propensity assay 1, except that none of the assayed worms were exposed to the parasite prior to the assay. See supplementary materials and (Slowinski et al.2023) for further details on the source of the nematode populations and on the protocols used in these assays.

### Calculating outcrossing rates

The proportion of offspring produced by outcrossing following mating opportunity on each mating plate was estimated based on offspring male frequencies (Stewart and Phillips 2002, Bahrami and Zhang 2013) using the following equations:

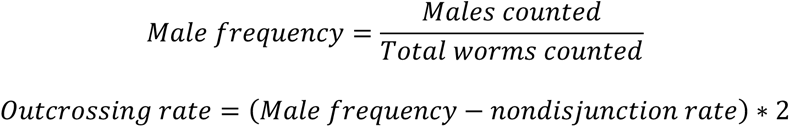

We assumed a nondisjunction rate of 0.0015 (Ward and Carrel 1979, Hodgkin and Doniach 1997). Note that while the true population outcrossing rates must be bounded by 0 and 1, our estimates of outcrossing rates are calculated based on male frequencies and are subject to sampling error. Hence it is possible for us to estimate an outcrossing rate greater than 1. For the purposes of our data presentation only, we constrained the plotted outcrossing rates to be less than one (i.e., we set outcrossing rates greater than one to one, and calculated mean and standard errors using the bounded outcrossing rates). However, for the purposes of our hypothesis-testing statistical analyses, we used our raw estimates of outcrossing rates. Note that if we run our statistics on the bounded outcrossing rates, we get qualitatively similar results (not shown).

### Determining sex of the mother in the evolution-of-outcrossing-propensity assays

Because the mothers in the evolution-of-outcrossing propensity were randomly selected from experimentally evolved trioecious populations that included both hermaphrodites and females, and because hermaphrodites and females cannot be easily distinguished morphologically, we identified the sex of the mothers in the evolution-of-outcrossing propensity assays using molecular methods (see supplementary materials). Outcrossing rates on mating plates with hermaphrodite mothers were analyzed separately from outcrossing rates on mating plates with female mothers.

### Outcrossing propensity assays excluded data

Two-worm protocol mating plates on which we could not find and remove both parents, or on which one or both parents had died during the 48-hour window of mating opportunity, or on which we were unable to unambiguously determine the sex of the mother, were excluded from our analyses. The hermaphrodites on some plates produced very few offspring. In such cases, if we were unable to score the sex of at least 20 offspring, the plate was excluded from our analyses. Very few plates were excluded due to insufficient offspring number (n = 4 out of 98 plates in the evolution-of-outcrossing-propensity assays, n = 6 out of 58 plates in the plasticity-of-outcrossing-propensity assay 1, n = 20 out of 230 plates in the plasticity-of-outcrossing-propensity assay 2) or due to one or both parents dead or missing at the end of the window of mating opportunity (n = 0 in the evolution-of-outcrossing-propensity assay, n = 2 in the plasticity-of-outcrossing-propensity assay 1, n = 0 in the plasticity-of-outcrossing propensity assay 2).

### Assays to determine whether outcrossing affects offspring resistance against the coevolving parasite

To test whether outcrossing affects offspring resistance against the coevolving parasite, individual hermaphrodites were isolated from host populations that had previously coevolved with the *S. marcescens* parasites for 24 host generations (Slowinski et al. 2023). Each hermaphrodite was then selfed for eight generations to create a highly homozygous lineage of hermaphrodites derived from a single genotype (hereafter referred to as a hermaphrodite lineage). Then hermaphrodites from each lineage were either selfed or outcrossed with randomly selected males from their source population. The offspring of these matings were then selfed for one more generation to produce F2 offspring that were then exposed to the *S. marcescens* parasite. The reason that we assayed the parasite resistance of the F2 offspring, rather than the F1 offspring following the selfing or outcrossing treatment, was to avoid confounding the outcrossed versus selfed status of the host offspring in our assays with sex. Because outcrossing produces a 1/1 ratio of males to hermaphrodites, while almost all selfed offspring are hermaphrodites, about half of the F1 offspring in our outcrossed treatment were males while almost all of the F1 offspring in our selfed treatment were hermaphrodites. The F1 males in the outcrossed treatment could have impaired our ability to detect effects of outcrossing on host resistance, if we had compared resistance in the F1s, as *C. elegans* males have previously been shown to be more susceptible than hermaphrodites to *S. marcescens* (Lynch et al. 2018). On the other hand, because we selfed F1 hermaphrodites in both treatments to produce the F2 progeny, almost all of the F2 progeny from both treatments were hermaphrodites. Hence our assay of F2 offspring allowed us to compare the parasite resistance of hermaphrodites from lineages with a recent history of outcrossing, versus the parasite resistance of hermaphrodites from fully selfing lineages.

Experimental parasite exposures were conducted on *Serratia* selection plates (SSPs), which are Petri plates filled with NGM lite and seeded with *S. marcescens* on one side of the plate, and *E. coli* on the other side of the plate (previously described in Morran et al. 2011, Slowinski et al. 2016, Slowinski et al. 2023). SSPs were seeded with the sympatric coevolved *S. marcescens* population (from generation 24), and survival was assessed 48 hours after exposure.

### Statistical methods

All p-values reported are 2-tailed unless otherwise specified. Statistical tests were run using R version 4.3.1.

### Plasticity of outcrossing propensity in response to parasite exposure

In assay 1, all of the N2 outcrossing propensity data was collected on one day and all of CW1-30 outcrossing propensity data was collected on another day. We used a two-way ANOVA to test whether nematode population (CW1-30 or N2) and parasite exposure treatment (parasite-exposed or control) affected outcrossing rates on our mating plates. We treated nematode population and parasite exposure treatment as fixed factors, and outcrossing rate as our dependent variable.

In assay 2, outcrossing propensity was assayed in 6 different blocks of mating plates, with parasite exposure treatment balanced across blocks, and with each worm strain assayed in 1 to 3 blocks. We used a linear mixed model with worm strain and parasite exposure treatment, and with a worm strain by parasite exposure treatment interaction to predict male frequencies. We treated experimental block as a random effect.

### Evolution of outcrossing propensity Hermaphrodite outcrossing rates

We used a linear mixed-effects model (LMM) to test whether population history of (co)evolution with the parasite predicted outcrossing rates on mating plates. Because each replicate host population was the source of the parents of multiple mating plates, we included replicate population (nested within treatment) as a random effect.

### Sex ratios in the offspring of female mothers

As a validation of our methods for determining the sex of mothers in the evolution-of-outcrossing propensity assays, we used a one-sample t-test to assess whether the offspring sex ratio of nematodes that were determined to be females (based on our molecular sexing assays) differed significantly from the expected frequency of 50%. For this analysis we pooled mating plates across treatments and across population replicates. Our analysis confirmed that the male frequency of offspring on mating plates with mothers that we had identified as females did not differ significantly from 0.5, which is the frequency that we would expect if mothers were females and all of their offspring were outcrossed. The mean male frequency on plates with mothers identified as female was 0.508349 (95% CI: 0.484-0.533, t_30_ = 0.704, *P* = 0.487).

### Survival of selfed versus outcrossed offspring exposed to the coevolved parasite

To evaluate whether outcrossing affected F2 offspring survival rates on the coevolved parasite we used Generalized Linear Models (GLM) with hermaphrodite lineage and outcrossing treatment as fixed factors, and with proportion of worms that survived 48 hours of exposure on the coevolved parasite as our dependent variable. The host offspring in our survival assay were derived from five different hermaphrodite lineages; all five hermaphrodite lineages were isolated from the same experimentally evolved host population. We used a maximum likelihood estimation method, an identity link function, and we fit a normal distribution. Pearson tests did not detect significant levels of overdispersion in the survival rates. We tested for a treatment by hermaphrodite lineage interaction, for a main effect of hermaphrodite lineage, and for a main effect of outcrossing treatment on survival. We performed linear contrast tests within the GLM to compare the selfed versus outcrossed treatments within each host genotype to determine which genotypes showed a significant effect of outcrossing treatment on offspring survival.

## Results

### Plasticity of outcrossing propensity

In assay 1, in which two genetically distinct nematode host populations were exposed for two hours to either a *S. marcescens* parasite treatment or to an *E. coli* food source control treatment, we found no significant interaction between nematode population and parasite treatment on outcrossing rates (*F_1,48_* = 0.003, *P* = 0.959, Figure 1A). We also found no significant main effect of parasite exposure treatment on outcrossing rates (*F_1,48_* = 0.312, *P* = 0.579, Figure 1A). However, there was a statistically significant main effect of nematode population on outcrossing rates, with the genetically diverse population CW1-30 (derived from the Hawaiian strain CB4856) exhibiting higher outcrossing rates than the inbred lab strain N2 (*F_1,48_* = 119.1, *P* < 0.001, Figure 1A).

**Figure 1:**
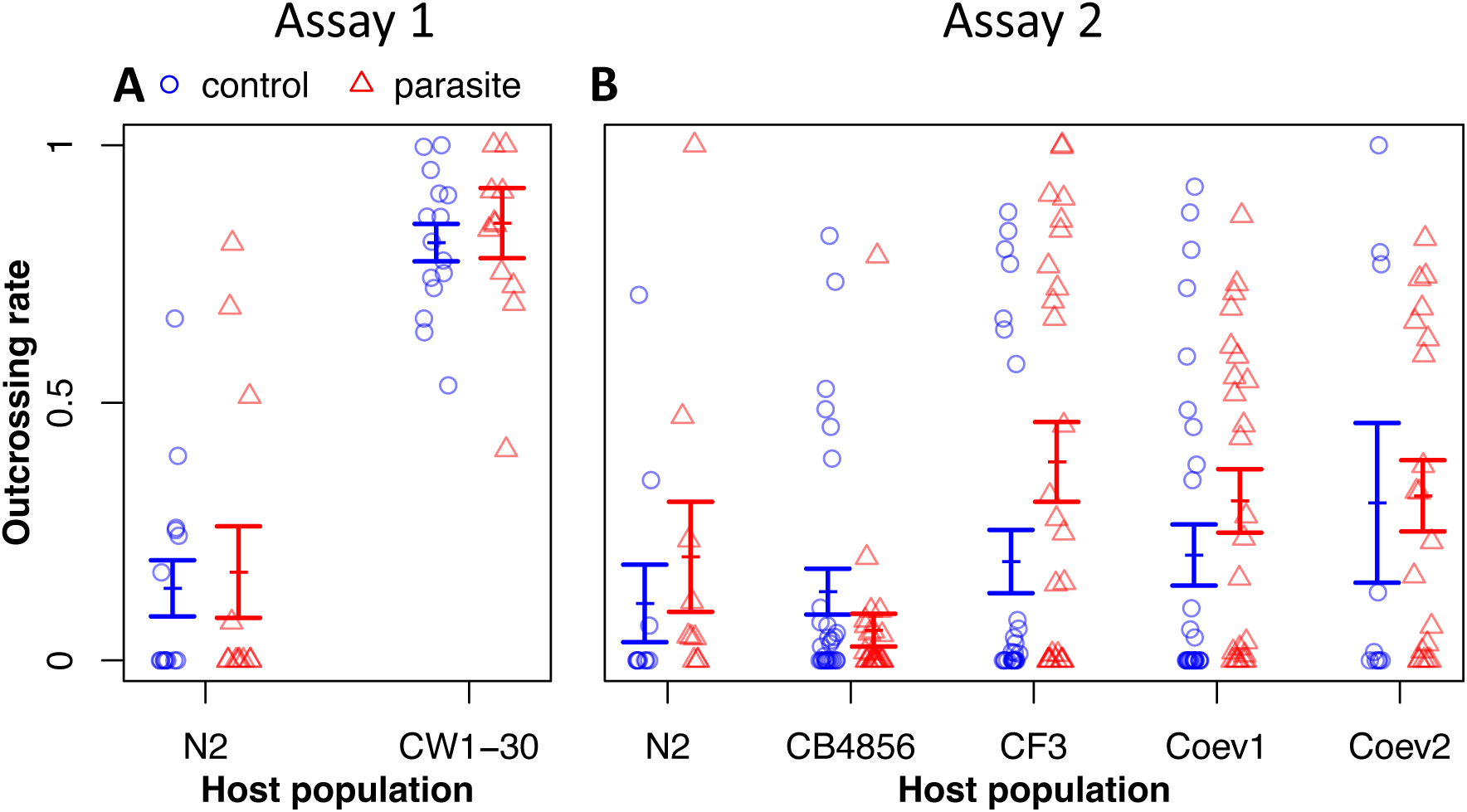
Outcrossing rates of worms following brief exposure to the *S. marcescens* parasite strain SM2170 (red triangles) or to a control, non-pathogenic *E. coli* food source (blue circles). N2 and CB4856 are inbred *C. elegans* lab strains. CW1-30 and CF3 are *C. elegans* populations that have been mutagenized and then passaged in the absence of the parasite. Coev1 and Coev2 represent genotypes isolated from a host population that had previously been copassaged with the *S. marcescens* pathogen for 24 host generations. Assay 1 (panel A): hermaphrodites were exposed for two hours and then one male and one hermaphrodite were paired on each mating plate. Assay 2 (panel B): hermaphrodites were exposed for four hours and then 10 hermaphrodites and 10 males were picked onto each mating plate. Each point represents the outcrossing rate observed on one mating plate. Error bars represent ± one standard error of the mean.

Similarly, in assay 2, in which five nematode host populations were exposed to a parasite or control treatment for four hours, we found no significant nematode-population by parasite treatment interaction on outcrossing rates (*F_4, 196_* = 1.7, *P* = 0.14, Figure 1B). We also found no significant main effect of parasite exposure on outcrossing rates (*F_1, 198_* = 1.24, *P* = 0.27). In contrast to assay 1, we found no significant main effect of nematode population on outcrossing rates in assay 2 (*F_4, 10_* = 2.1, *P* = 0.158, Figure 1B).

### Evolution of outcrossing propensity

In a control (parasite-free) assay environment, we compared hermaphrodite outcrossing rates among populations with different histories of (co)evolutionary interactions with the *S. marcescens* parasite. We found no effect of (co)evolutionary history with the parasite on outcrossing rates on mating plates with hermaphrodite mothers (LMM, *F_2_* = 0.07, *P* = 0.93, Figure 2).

**Figure 2:**
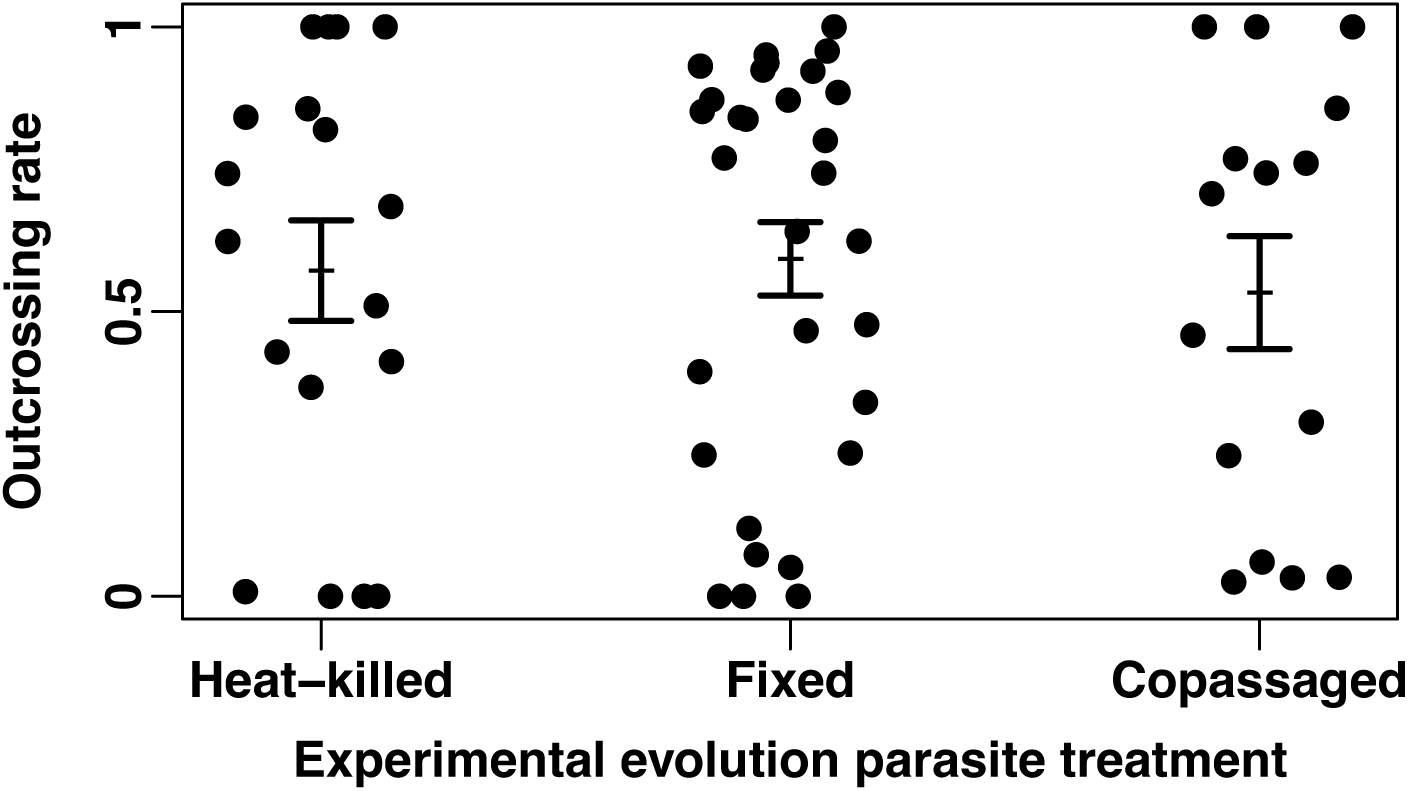
Outcrossing rates on mating plates with hermaphrodite mothers from populations that had previously evolved for 24 host generations in the heat-killed parasite treatment (left), the fixed parasite treatment (middle), or the copassaged parasite treatment (right). Each point represents the outcrossing rate observed on one mating plate. Error bars represent ± one standard error of the mean.

### Effect of hermaphrodite lineage and outcrossing treatment on the parasite resistance of F2 host offspring

We found a significant main effect of hermaphrodite lineage (GLM: *P* < 0.001, Figure 3a) and a significant main effect of outcrossing treatment (*P* < 0.001, Figure 3A and B) on survival of F2 host offspring on the parasite. We also found a significant hermaphrodite-genotype by outcrossing-treatment interaction on the survival of F2 hosts on the parasite (*P* < 0.001, Figure 3A). Linear contrasts revealed that outcrossing significantly increased offspring survival for hermaphrodite lineage 2 (Chi-Square = 6.58, *P* = 0.01), hermaphrodite lineage 3 (Chi-Square = 13.84, *P* < 0.001), and hermaphrodite lineage 4 (Chi-Square = 27.65, *P* < 0.001) (Figure 3A).

**Figure 3:**
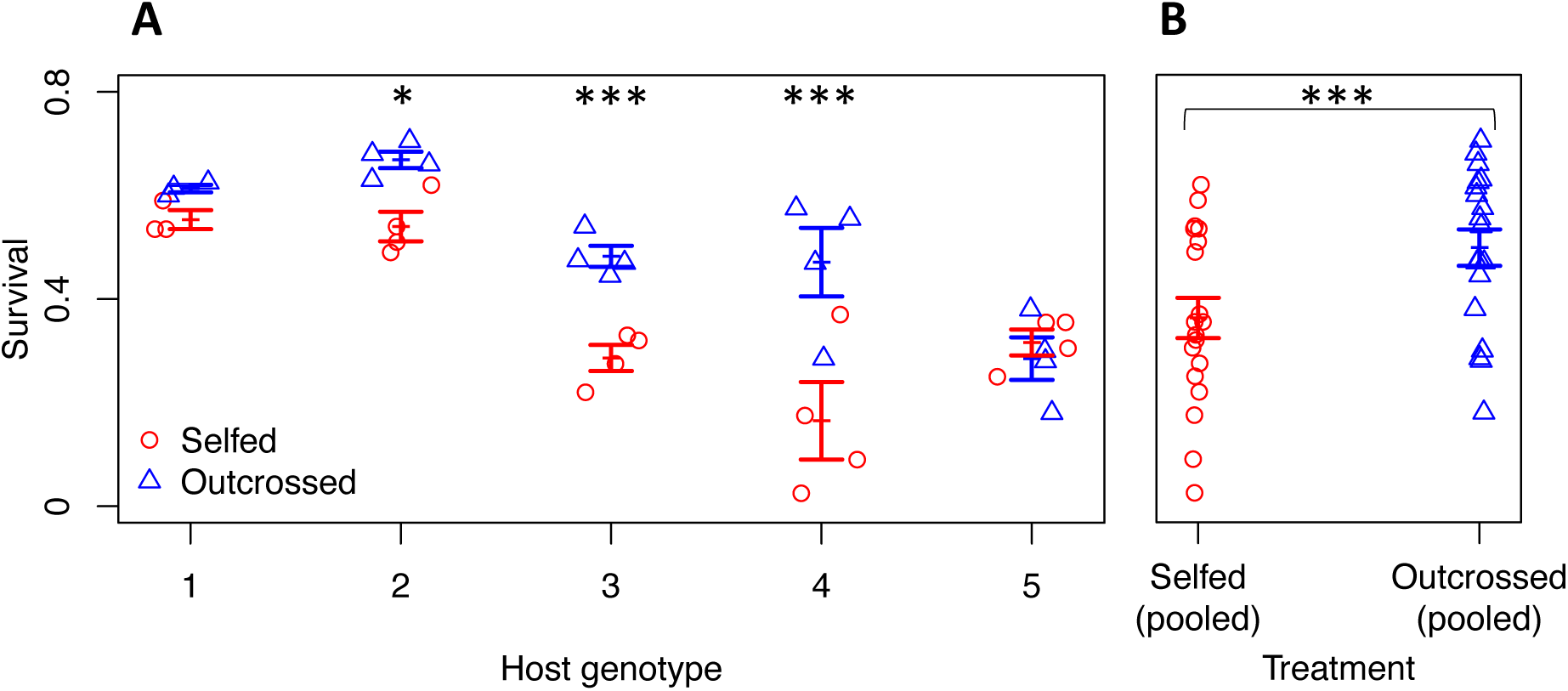
Survival of selfed (red circle) and outcrossed (blue triangle) F2 *C. elegans* offspring following 48 hours of exposure to their coevolved *S. marcescens* parasite. Points represent the proportion of offspring that survived parasite exposure on each replicate assay plate. All hermaphrodite lineages in this assay were isolated from the same replicate host population. Horizontal lines represent the mean proportion of offspring that survived 48 hours of parasite exposure across outcrossed or selfed replicates. Error bars represent ± 1 standard error of the mean. A) Survival rates for offspring of each hermaphrodite lineage are presented separately. B) Offspring pooled within treatments across hermaphrodite lineages. * *P* < 0.05; *** *P* < 0.001

## Discussion

Previous research has demonstrated that coevolutionary interactions with parasites can maintain high outcrossing rates in *C. elegans* host populations (Morran et al. 2011, Slowinski et al. 2016, Slowinski et al. 2023). We tested whether parasite-induced changes in host outcrossing propensity could explain how coevolving parasites maintain high host outcrossing rates. We found no effect of brief (within-generation) exposure to the bacterial parasite *S. marcescens* on *C. elegans* outcrossing propensity, suggesting that *C. elegans* do not plastically increase their propensity to outcross in response to exposure to parasites in their environment. Furthermore, we found no effect of prior evolution or coevolution with the parasite on outcrossing propensity, suggesting that *C. elegans* outcrossing propensity did not evolve in response to selective pressure from a coevolving parasite, at least not on the evolutionary timescale of our experiment (24 host generations). Next, we tested whether selective culling of selfed offspring by parasites from host populations could explain how coevolving parasites maintain high host outcrossing rates. We found that parental outcrossing significantly increased the resistance of F2 offspring against a coevolving parasite. Taken together these results suggest that coevolving parasites maintain high outcrossing rates in *C. elegans* host populations by selecting against selfed offspring, and not by causing changes in host outcrossing propensity.

### Plasticity of outcrossing propensity

We found no effect of brief (within-generation) exposure to *S. marcescens* on outcrossing propensity in *C. elegans*. This result contrasts with recent research in other host-parasite study systems showing that hosts can plastically diversify their offspring in response to exposure to parasites (Soper et al. 2014, Singh et al. 2015, Hite et al. 2017). Furthermore, our results contrast with recent research (Wu et al. 2023) in our own host system, showing that exposure to the pathogenic bacteria *P. aeruginosa* can induce increased outcrossing propensity in *C. elegans* hosts. Hence, the ability to induce increased *C. elegans* outcrossing propensity may be pathogen-specific, with *P. aeruginosa* able to induce changes while *S. marcescens* cannot.

Alternatively, it is possible that *S. marcescens* can induce plastic changes in *C. elegans* outcrossing propensity, leading to higher offspring diversity, but that the parasite-exposure protocol in our study was not sufficient to induce this effect. We exposed *C. elegans* to the *S. marcescens* parasite for about two hours (assay 1) or four hours (assay 2). Perhaps a longer exposure to *S. marcescens* would be necessary to induce plastic changes in host outcrossing propensity. However, we consider this possibility to be unlikely, because in another study (Kurz et al. 2003), two hours of exposure of L4 *C. elegans* to the GFP-tagged *S. marcescens* strain Db11 was sufficient for intact bacteria to be observed accumulating in the lumen of the intestine, suggesting that our treatment was likely a significant immunological challenge. Additionally, Mallo et al. (2002) demonstrated that exposure to the *S. marcescens* strain Db11 for less than 6 hours was sufficient for intact *S. marcescens* bacteria to get into the host intestinal lumen and start proliferating, and also to induce upregulation of host genes associated with immune defense. Furthermore, the *S. marcescens* parasite strain in our assays (Sm2170) is so virulent that a longer exposure would significantly increase the probability of host mortality. Finally, four hours of exposure to *P. aeruginosa* (the same duration that we exposed *C. elegans* to *S. marcescens* in assay 2) was recently shown to induce increased outcrossing propensity in *C. elegans* (Wu et al. 2023). Regardless of whether or not a longer parasite exposure could induce changes in host outcrossing propensity, plastically induced increases in host outcrossing propensity are unlikely to explain how parasites maintained the high outcrossing rates previously reported in *C. elegans* experimental evolution studies (Morran et al. 2011, Slowinski et al. 2016, Slowinski et al. 2023); the 2-4 hour duration of exposure in our outcrossing propensity assays was approximately consistent with the duration of exposure the hosts in those studies typically experienced before moving off of the lawn of the *S. marcescens* parasite (LTM personal observation).

### Evolution of outcrossing propensity

We found no difference in the outcrossing propensity of hermaphrodites sampled from populations that had previously evolved in the presence or absence of a coevolving parasite. This result suggests that *C. elegans* outcrossing propensity does not evolve in response to evolutionary, or coevolutionary, interactions with parasites, at least not on the time scale of our experiment (24 host generations). One possible explanation for why outcrossing propensity did not evolve is that coevolution with the parasite did not produce strong selection on outcrossing propensity. However, we consider this explanation to be unlikely. Morran et al. (2011), Slowinski et al. (2016), and Slowinski et al. (2023) found that *C. elegans* host populations coevolving with the *S. marcescens* parasite maintained high outcrossing rates relative to host populations not coevolving with the parasite, suggesting that parasite-mediated selection for outcrossing was sufficiently strong to offset the two-fold cost of males. Furthermore, in the present study, parental outcrossing increased the resistance of F2 host offspring exposed to the coevolving parasite. These results strongly suggest that the coevolving parasite favored outcrossing and would be expected to select for host genotypes that preferentially reproduce by outcrossing as opposed to self-fertilization. Still, selection for outcrossing may have been weaker than selection on loci that directly mediated parasite resistance. An alternative possible explanation for why were unable to detect evolution of outcrossing propensity is that there was little or no additive genetic variance for outcrossing propensity in the host populations in our experiment. In this scenario, the evolution of outcrossing propensity would be constrained even if coevolution with the parasite favored increased outcrossing propensity.

### Parasite resistance of selfed versus outcrossed offspring

We found that outcrossing in the parental generation significantly increased the resistance of F2 host offspring exposed to the coevolving parasite. This result suggests that, in an environment with a coevolving parasite, parasites select against selfing host lineages. Hence outcrossing benefits hosts by conferring protection against parasites for their offspring. Because outcrossing produces a 1/1 ratio of males to hermaphrodites while almost all offspring produced by self-fertilization are hermaphrodites, the selective culling of selfed offspring that we observed would primarily remove selfed hermaphrodites from host populations coevolving with a parasite, increasing host population male frequencies. This would increase the opportunity for outcrossing in host populations, and hence could explain how parasites maintain high outcrossing rates in coevolving host populations, even in the absence of changes in host outcrossing propensity.

## Conclusions

The Red Queen hypothesis has received broad empirical support, including several recent experimental evolution studies demonstrating that coevolutionary interactions with the virulent bacterial parasite *Serratia marcescens* can maintain high outcrossing rates in the nematode host *Caenorhabditis elegans* (Morran et al. 2011, Slowinski et al. 2016, Slowinski et al. 2023). In the present study, we found no evidence that *S. marcescens* parasites induce plastic or evolved changes in the outcrossing propensity of *C. elegans* hosts, suggesting that parasites are unlikely to maintain high host outcrossing rates by altering the outcrossing propensity of their hosts. We found that parental outcrossing increased the parasite resistance of host offspring in the F2 generation. This suggests that coevolving parasites select against selfing host lineages and maintain high outcrossing rates by selectively culling selfed offspring (mostly hermaphrodites) from host populations.

## Author contributions

SPS, LTM, and CML conceptualized the original project and designed the experiments. SPS ran the plasticity of outcrossing propensity assay 1, and the evolution of outcrossing propensity assay. JDG ran the plasticity of outcrossing propensity assay 2. MJP ran the survival assays of selfed versus outcrossed offspring. LTM supervised all of the assays. SPS wrote the manuscript with editing help from LTM and CML. All authors read and approved the manuscript.

## Acknowledgements

We thank Eric Cui and Katya Haspel for help preparing media and seeding plates to maintain the nematodes in our assays. We thank Eric Cui and Amrita Bhattacharya for helping to develop protocols. We thank members of the Lively and Bashey labs and Laura Alexander for feedback on our study design and data analysis.

## Data availability

Data will be made available on Dryad upon acceptance of the manuscript.

## Literature Cited

Acevedo-Whitehouse, K., F. Gulland, D. Greig, and W. Amos. 2003. Disease susceptibility in California sea lions. Nature 422:35–35.

Bahrami, A. K., and Y. Zhang. 2013. When females produce sperm: genetics of *C. elegans* hermaphrodite reproductive choice. G3-Genes Genomes Genetics 3:1851–1859.

Bell, G. 1982. The masterpiece of nature: the evolution and genetics of sexuality. University of California Press, Berkeley.

Brenner, S. 1974. Genetics of *Caenorhabditis elegans*. Genetics 77:71–94.

Clay, K., and P. Kover. 1996. Evolution and stasis in plant-pathogen associations. Ecology 77:997–1003.

Duncan, A. B., and T. J. Little. 2007. Parasite-driven genetic change in a natural population of *Daphnia*. Evolution 61:796–803.

Dybdahl, M. F., and C. M. Lively. 1998. Host-parasite coevolution: Evidence for rare advantage and time-lagged selection in a natural population. Evolution 52:1057–1066.

Ellison, A., J. Cable, and S. Consuegra. 2011. Best of both worlds? Association between outcrossing and parasite loads in a selfing fish. Evolution 65:3021–3026.

Gibson, A. K., L. F. Delph, and C. M. Lively. 2017. The two-fold cost of sex: Experimental evidence from a natural system. Evolution Letters 1:6–15.

Hamilton, W. D. 1980. Sex versus non-sex versus parasite. Oikos 35:282–290.

Hite, J. L., R. M. Penczykowski, M. S. Shocket, K. A. Griebel, A. T. Strauss, M. A. Duffy, C. E. Cáceres, and S. R. Hall. 2017. Allocation, not male resistance, increases male frequency during epidemics: a case study in facultatively sexual hosts. Ecology 98:2773–2783.

Hodgkin, J., and T. Doniach. 1997. Natural variation and copulatory plug formation in *Caenorhabditis elegans*. Genetics 146:149–164.

Jaenike, J. 1978. An hypothesis to account for the maintenance of sex within populations. Evolutionary Theory 3:191–194.

Jokela, J., M. F. Dybdahl, and C. M. Lively. 2009. The maintenance of sex, clonal dynamics, and host-parasite coevolution in a mixed population of sexual and asexual snails. American Naturalist 174:S43–S53.

Jokela, J., and C. M. Lively. 1995. Spatial variation in infection by digenetic trematodes in a population of freshwater snails (*Potamopyrgus antipodarum*). Oecologia 103:509–517.

Keller, L. F., and D. M. Waller. 2002. Inbreeding effects in wild populations. Trends in Ecology & Evolution 17:230–241.

King, K. C., J. Jokela, and C. M. Lively. 2011. Parasites, sex, and clonal diversity in natural snail populations. Evolution 65:1474–1481.

King, K. C., and C. M. Lively. 2009. Geographic variation in sterilizing parasite species and the Red Queen. Oikos 118:1416–1420.

King, K. C., and C. M. Lively. 2012. Does genetic diversity limit disease spread in natural host populations? Heredity 109:199–203.

Krist, A. C., C. M. Lively, E. P. Levri, and J. Jokela. 2000. Spatial variation in susceptibility to infection in a snail-trematode interaction. Parasitology 121:395–401.

Kurz, C. L., S. Chauvet, E. Andres, M. Aurouze, I. Vallet, G. P. F. Michel, M. Uh, J. Celli, A. Filloux, S. de Bentzmann, I. Steinmetz, J. A. Hoffmann, B. B. Finlay, J. P. Gorvel, D. Ferrandon, and J. J. Ewbank. 2003. Virulence factors of the human opportunistic pathogen *Serratia marcescens* identified by in vivo screening. Embo Journal 22:1451–1460.

Lively, C. M. 1987. Evidence from a New Zealand snail for the maintenance of sex by parasitism. Nature 328:519–521.

Lively, C. M. 1992. Parthenogenesis in a freshwater snail: reproductive assurance versus parasitic release. Evolution 46:907–913.

Lively, C. M., and M. F. Dybdahl. 2000. Parasite adaptation to locally common host genotypes. Nature 405:679–681.

Lively, C. M., and J. Jokela. 1996. Clinal variation for local adaptation in a host-parasite interaction. Proceedings of the Royal Society B-Biological Sciences 263:891–897.

Lively, C. M., and J. Jokela. 2002. Temporal and spatial distributions of parasites and sex in a freshwater snail. Evolutionary Ecology Research 4:219–226.

Lynch, Z. R., M. J. Penley, and L. T. Morran. 2018. Turnover in local parasite populations temporarily favors host outcrossing over self-fertilization during experimental evolution. Ecol Evol 8:6652–6662.

Mallo, G. V., C. L. Kurz, C. Couillault, N. Pujol, S. Granjeaud, Y. Kohara, and J. J. Ewbank. 2002. Inducible antibacterial defense system in C-elegans. Current Biology 12:1209–1214.

Masri, L., R. D. Schulte, N. Timmermeyer, S. Thanisch, L. L. Crummenerl, G. Jansen, N. K. Michiels, and H. Schulenburg. 2013. Sex differences in host defence interfere with parasite-mediated selection for outcrossing during host-parasite coevolution. Ecology Letters 16:461–468.

Maynard Smith, J. 1978. The evolution of sex. New York, Cambridge [Eng.].

Morran, L. T., B. J. Cappy, J. L. Anderson, and P. C. Phillips. 2009. Sexual partners for the stressed: facultative outcrossing in the self-fertilizing nematode *Caenorhabditis elegans*. Evolution 63:1473–1482.

Morran, L. T., O. G. Schmidt, I. A. Gelarden, R. C. Parrish, II, and C. M. Lively. 2011. Running with the Red Queen: host-parasite coevolution selects for biparental sex. Science 333:216–218.

Parrish, R. C., M. J. Penley, and L. T. Morran. 2016. The integral role of genetic variation in the evolution of outcrossing in the *Caenorhabditis elegans*-*Serratia marcescens* host-parasite system. PLOS ONE 11.

Singh, N. D., D. R. Criscoe, S. Skolfield, K. P. Kohl, E. S. Keebaugh, and T. A. Schlenke. 2015. Fruit flies diversify their offspring in response to parasite infection. Science 349:747–750.

Slowinski, S. P., J. Cho, M. J. Penley, L. W. Alexander, A. B. Greenberg, S. R. Namburar, and L. T. Morran. 2023. High parasite virulence necessary for the maintenance of host outcrossing via parasite-mediated selection. Evolution Letters:qrad036.

Slowinski, S. P., L. T. Morran, R. C. Parrish, E. R. Cui, A. Bhattacharya, C. M. Lively, and P. C. Phillips. 2016. Coevolutionary interactions with parasites constrain the spread of self-fertilization into outcrossing host populations. Evolution 70:2632–2639.

Soper, D. M., K. C. King, D. Vergara, and C. M. Lively. 2014. Exposure to parasites increases promiscuity in a freshwater snail. Biology letters 10:20131091–20131091.

Spielman, D., B. W. Brook, D. A. Briscoe, and R. Frankham. 2004. Does inbreeding and loss of genetic diversity decrease disease resistance? Conservation Genetics 5:439–448.

Stewart, A. D., and P. C. Phillips. 2002. Selection and maintenance of androdioecy in *Caenorhabditis elegans*. Genetics 160:975–982.

Verhoeven, K. J. F., and A. Biere. 2013. Geographic parthenogenesis and plant-enemy interactions in the common dandelion. BMC Evolutionary Biology 13:23–32.

Vrijenhoek, R. C. 1998. Animal clones and diversity. Bioscience 48:617–628.

Ward, S., and J. S. Carrel. 1979. Fertilization and sperm competition in the nematode *Caenorhabditis elegans*. Dev Biol 73:304–321.

Wolinska, J., and P. Spaak. 2009. The cost of being common: evidence from natural *Daphnia* populations. Evolution 63:1893–1901.

Wu, T., M. Ge, M. Wu, F. Duan, J. Liang, M. Chen, X. Gracida, H. Liu, W. Yang, A. R. Dar, C. Li, R. A. Butcher, A. L. Saltzman, and Y. Zhang. 2023. Pathogenic bacteria modulate pheromone response to promote mating. Nature 613:324–331.

