## Supplemental material for "Outcrossing increases resistance against coevolving parasites"

**Supplemental material for the manuscript "Outcrossing increases resistance against coevolving parasites"**

*Synchronizing lines for outcrossing propensity assays*

Prior to outcrossing propensity assays, the life-stages of the nematodes in assay populations were synchronized using a "hatch off" protocol (Stiernagle 2006). In the hatch off, nematode populations were treated with 120 µl of 60% bleach solution mixed into 1,000 µl of a solution of nematodes suspended in M9 to kill all the nematode life stages except the eggs (Stiernagle 2006). Following exposure to the bleach solution, nematodes were washed with sterile deionized water and then continuously mixed in an M9 solution for 24 hours in a tube rotator, during which period the eggs hatched into L1 larvae. L1 larvae were transferred onto plates that had been constructed by pouring 24 mL of autoclaved nematode growth medium (NGM) Lite (US Biological, Swampscott, MA) onto a 10 cm Petri dish. The NGM had been seeded with a lawn of 60 µl of the *E. coli* strain OP50, which had grown overnight in a 28 °C incubator before the nematodes were transferred onto the plate. After the L1 nematodes were transferred onto the seeded OP50 plate they grew at 20 °C for 48 hours until they matured.

*Source of the nematode lines used in the plasticity-of-outcrossing-propensity assay 1*

Nematode populations were derived from the nematode strain CB4856, originally from Hawaii, which was provided by the Caenorhabditis Genetics Center (University of Minnesota, Minneapolis, MN). Prior to our experiment, a line called PX382 was derived by systematically inbreeding the strain CB4856 (Morran et al. 2009). An obligate outcrossing strain called PX386 was derived from the inbred strain by systematically backcrossing the mutant allele *fog-2(q71)* into the genetic background of PX382 (Morran et al. 2009). Prior to our experiment, during three consecutive generations, five near-isogenic populations of PX386 were independently mutagenized at 40mM of EMS for four hours to create five genetically diverse obligately outcrossing populations (fixed for the obligate-outcrossing allele (*fog-2(q71)*). Five near-isogenic populations of PX382 were also mutagenized (same protocol) to create five genetically diverse mixed-mating populations (fixed for the mixed-mating allele (*fog-2(wt)*) (Morran et al. 2011). This procedure induced approximately 1,000 point mutations per lineage in each population (Epstein and Shakes 1995, Morran et al. 2011). The genetically diverse mixed-mating and obligately outcrossing nematode populations were passaged for 30 generations under 3 experimental treatments (Morran et al. 2011). One of the genetically diverse mixed-mating populations (called CW1-30) that had been passaged for 30 generations under the heat-killed parasite treatment in the (Morran et al. 2011) study was used as the source of hermaphrodites on the mating plates. One of the genetically diverse obligately outcrossing populations (designated F5) that was frozen prior to the (Morran et al. 2011) study (i.e. a population of obligately outcrossing individuals that was an ancestor to the populations in the (Morran et al. 2011) study) was used as the source of males on the mating plates. We also assayed the outcrossing propensity of the inbred lab strain N2 (provided by the Caenorhabditis Genetics Center, originally isolated from Britain).

*Source of the nematode lines used in the evolution-of-outcrossing-propensity assays, the resistance of selfed versus outcrossed offspring assays, and in the plasticity-of-outcrossing-propensity assay 2*

Prior to the present study, the mixed-mating allele from the PX382 strain (see above) was introgressed through a series of seven backcrosses into the genetic background of 12 obligately outcrossing populations from the (Morran et al. 2011) study to create 12 mixed-mating populations each representing a genetic background derived from one of the obligately outcrossing populations (Slowinski et al. 2016). Slowinski et al. (2023) combined nematodes from one of those obligately outcrossing populations (designated CF3, which had evolved under the heat-killed parasite treatment in the (Morran et al. 2011) study), with nematodes from its associated mixed-mating strain (i.e. with nematodes bearing the mixed mating allele and the same genetic background), at a starting frequency of 10% mixed-maters, to create a trioecious experimental population (i.e., composed of males, females, and hermaphrodites). The trioecious population was divided into four replicate populations, and each of the four replicate populations was divided into three experimental treatments, which were passaged on *Serratia* selection plates (SSPs) for 24 host *C. elegans* generations (Slowinski et al. 2023). Briefly, nematode populations in the heat-killed parasite treatment were passaged on SSPs seeded with heat-killed (avirulent) SM2170 *Serratia* strain. Nematode populations in the fixed parasite treatment were passaged on SSPs seeded with a stock, virulent, but non-evolving population of SM2170. Nematode populations in the copassaged treatment were copassaged with an evolving (and potentially coevolving) strain of SM2170, which was originally seeded from the same stock population used in the fixed parasite treatment, but which evolved over the course of the study.

After 24 generations of experimental evolution in (Slowinski et al. 2023), the evolved nematode strains (from the heat-killed parasite treatment, the fixed parasite treatment, and the copassaged parasite treatment) were assayed (in the present study) in a control (no-parasite) environment to determine whether outcrossing propensity evolved in response to (co)evolutionary interactions with the parasite (see below for outcrossing propensity assay details). Additionally, five hermaphrodite genotypes were isolated from one of the copassaged host populations, inbred, and the survival of their selfed versus outcrossed offspring on their contemporary copassaged parasite was measured. Finally, the outcrossing propensity of two of the inbred hermaphrodite genotypes from a copassaged host population was assayed in the plasticity-of-outcrossing-propensity assay 2.

A schematic showing the source of the nematode populations is provided (Figure S1).

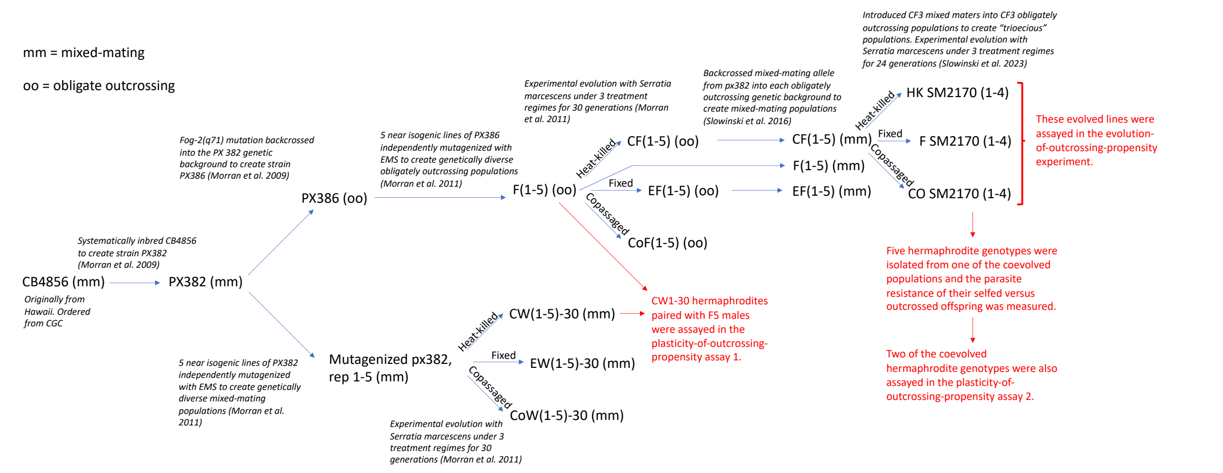


Figure S1: A schematic showing the source of the nematode populations assayed in the present study.

*Evolution-of-outcrossing-propensity experiment assay details*

Following 24 generations of experimental evolution with the *S. marcescens* strain SM2170 in either the heat-killed parasite, the fixed parasite, or the copassaged parasite treatment (Slowinski et al. 2023), experimental nematode populations were maintained at 15 °C and transferred to fresh OP50-seeded plates once per week until their outcrossing propensity was assayed. At 15 °C, the *C. elegans* life cycle is completed in about one generation per six or seven days (Byerly et al. 1976). Outcrossing propensity was assayed four to nine weeks (estimated four to nine generations) after the experimental lines were removed from selection in their respective experimental evolution regimes. The amount of time that experimental populations were maintained out of selection prior to being assayed did not differ across treatments. We confirmed that our experimental populations had maintained treatment differences in male frequencies during the period in which they were maintained out of selection prior to our outcrossing propensity assays.

Outcrossing propensity was assayed on 10 replicate mating plates from each of the four replicate heat-killed parasite treatment populations and from each of the four replicate fixed-parasite treatment host populations. Outcrossing propensity was assayed on 20 replicate mating plates from each of the four replicate copassaged host populations. The populations assayed in this experiment had evolved from trioecious starting populations (i.e. populations composed of hermaphrodites, females, and males), which segregated both the mixed-mating allele *fog-2(wt)* and the obligate-outcrossing allele *fog-2(q71)*. Most of these populations were still composed of all three sexes (hermaphrodites, females, and males) and still segregated both the mixed-mating allele and the obligate-outcrossing allele when we assayed them, although some of the populations might have lost one or the other *fog-2* alleles over the course of experimental evolution. When picking mating plates for these assays we paired each L4 male with a partner who we determined based on morphology to be either a hermaphrodite or a female. Because females (i.e., individuals with two X chromosomes that are homozygous for the obligate-outcrossing allele) must outcross with males to reproduce, whereas hermaphrodites (i.e., individuals with two X chromosomes that express the mixed-mating allele) can reproduce by either outcrossing or by self-fertilization, it was critical for us to distinguish between hermaphrodite mothers and female mothers in the analysis of our outcrossing propensity assay results. Hermaphrodites cannot be easily distinguished morphologically from females. We used molecular methods (see below) to determine the genotype of the mothers at the *fog-2* locus, which allowed us to determine the sex of the mothers in our outcrossing propensity assays. When we removed the parents from the mating plates after 48 hours of mating opportunity, we picked each mother into a separate proteinase K solution and her DNA was extracted for *fog-2* genotyping assays.

*Genotyping assays to determine the sex of the mother on each mating plate (evolution-of-outcrossing-propensity experiment)*

We used polymerase chain reactions (PCRs), in a separate reaction for each individual mother, to amplify the *fog-2* locus to determine the *fog-2* genotype of the mother from each of our mating plates. DNA was extracted when the parents were removed from the mating plates (48 hours after they were picked as L4 larvae onto the mating plates) by picking the nematodes into a proteinase K solution and thermal-cycling the solution. Extracted DNA was stored at -20 °C until it was used in the PCRs. We used the published PCR primers F1RFLP, FogR4short, and FogR3 (Theologidis et al. 2014). We validated our results by running positive controls (i.e., a nematode with known homozygous *fog-2(wt, wt)* genotype and a nematode with known homozygous *fog-2(q71, q71)* genotype) on every gel. Nematodes with a *fog-2(wt, wt)* genotype yield a 295 bp DNA band that can be visualized on agarose gels, while *fog-(q71, q71)* genotypes yield a 264 bp band (Theologidis et al. 2014, Slowinski et al. 2016). Previous studies have reported conflicting results for genotyping heterozygous *fog-2(wt, q71)* individuals using these primers. Theologidis et al. (2014) reported that *fog-2(wt, q71)* heterozygotes exhibited a double band pattern, while Slowinski et al. (2016) reported that *fog-2(wt, q71)* heterozygotes yielded only one visible DNA band (295 bp) and therefore were indistinguishable from *fog-2(wt, wt)* individuals in the assays. Because the goal of our PCR assays was to determine the sex (hermaphrodite or female) of the mother on each mating plate, it was not important for us to distinguish between *fog-2(wt, q71)* heterozygous genotypes and *fog-2(wt, wt)* homozygous genotypes because both genotypes will express a hermaphrodite phenotype. Therefore, in our assays, we scored individuals as hermaphrodites whenever we observed at least one DNA band at 295 bp, and we scored individuals as females whenever we observed only one band at 264 bp. Gels were scored treatment blind by SPS. As additional controls, we ran the PCRs and gels on known females *(fog-2(q71, q71))* that had mated with mixed-mating males *(fog-2(wt, wt))* and on known hermaphrodites *(fog-2(wt, wt))* that had mated with obligately outcrossing males *(fog-2(q71, q71))* to confirm that the number of gene copies provided by the male's sperm inside mated mothers is insufficient to compete for the amplification, such that it does not alter the apparent genotype of the mother in our genotyping assays.

**Literature cited**


Byerly, L., R. C. Cassada, and R. L. Russell. 1976. The life cycle of the nematode *Caenorhabditis elegans*: I. Wild-type growth and reproduction. Dev Biol **51**:23-33.

Epstein, H. F., and D. C. Shakes. 1995. *Caenorhabditis elegans*: modern biological analysis of an organism. Pages 31-54. Academic Press, San Diego.

Morran, L. T., M. D. Parmenter, and P. C. Phillips. 2009. Mutation load and rapid adaptation favour outcrossing over self-fertilization. Nature **462**:350-352.

Morran, L. T., O. G. Schmidt, I. A. Gelarden, R. C. Parrish, II, and C. M. Lively. 2011. Running with the Red Queen: host-parasite coevolution selects for biparental sex. Science **333**:216-218.

Slowinski, S. P., J. Cho, M. J. Penley, L. W. Alexander, A. B. Greenberg, S. R. Namburar, and L. T. Morran. 2023. High parasite virulence necessary for the maintenance of host outcrossing via parasite-mediated selection. Evolution Letters:qrad036.

Slowinski, S. P., L. T. Morran, R. C. Parrish, E. R. Cui, A. Bhattacharya, C. M. Lively, and P. C. Phillips. 2016. Coevolutionary interactions with parasites constrain the spread of self-fertilization into outcrossing host populations. Evolution **70**:2632-2639.

Stiernagle, T. 2006. Maintenance of *C. elegans*. Wormbook, ed. The C. elegans Research Community.

Theologidis, I., I. M. Chelo, C. Goy, and H. Teotonio. 2014. Reproductive assurance drives transitions to self-fertilization in experimental *Caenorhabditis elegans*. Bmc Biology **12**:93-114.
